# B Cell Receptor-Responsive miR-141 and viral miR-BART9 promote Epstein-Barr Virus Reactivation Through FOXO3 Inhibition

**DOI:** 10.1101/837294

**Authors:** Yan Chen, Devin Fachko, Nikita S. Ivanov, Rebecca L. Skalsky

**Affiliations:** Vaccine and Gene Therapy Institute, Oregon Health & Science University, Beaverton, Oregon

## Abstract

Antigen recognition by the B cell receptor (BCR) is a physiological trigger for reactivation of Epstein-Barr virus (EBV) and can be recapitulated in vitro by cross-linking of surface immunoglobulins. Previously, we identified a subset of EBV microRNAs (miRNAs) that attenuate BCR signal transduction and subsequently, dampen lytic replication in B cells. The roles of host miRNAs in virus reactivation are not completely understood. To investigate this process, we profiled the small RNAs in latently infected and reactivated Burkitt’s lymphoma cells, and identified several miRNAs, such as miR-141, that are induced upon BCR cross-linking. Notably, EBV encodes a viral miRNA, miR-BART9, with sequence homology to miR-141. To better understand the functions of these two miRNAs, we examined their molecular targets and experimentally validated multiple candidates commonly regulated by both miRNAs. Targets included transcriptional regulators of the EBV immediate early promoters and B cell transcription factors, leading us to hypothesize that these miRNAs modulate kinetics of the latent to lytic switch in B cells. Through functional assays, we identified roles for miR-141 and EBV miR-BART9 and one specific target, FOXO3, in lytic reactivation. Our data support a model whereby EBV exploits BCR-responsive miR-141 and further mimics activity of this miRNA family via a viral miRNA to promote productive virus replication.

**Importance:** EBV is a human pathogen associated with several malignancies. A key aspect of lifelong virus persistence is the ability to switch between latent and lytic replication modes. Mechanisms governing latency and lytic reactivation are only partly understood, and how the EBV latent to lytic switch is post-transcriptionally regulated remains an outstanding question. This study sheds light on how miR-141 expression is regulated in Burkitt’s lymphoma cells, and identifies a role for FOXO3, a common target of both miR-141 and viral miR-BART9, in modulating EBV reactivation.

## Introduction

Epstein-Barr virus (EBV) infects nearly 95% of adults worldwide. The virus primarily causes infectious mononucleosis (IM) in adolescents, is closely associated with lymphoproliferative disease in immunocompromised individuals, and is linked to cancers including Burkitt lymphomas (BL), diffuse large B cell lymphomas (DLBCL), and nasopharyngeal carcinoma (1, 2). BL is a common childhood cancer in Africa, and >80% of cases are EBV positive (3). As a germinal-center (GC) derived cancer, BL cells exhibit a centroblast-like phenotype (2, 4), maintained in part from activated c-myc oncogene expression (5), and have a molecular profile akin to GC dark zone proliferative B cells (6).

EBV has both a latent and a lytic replication phase, and periodically, latent EBV reactivates to produce infectious virions. The EBV lytic stage is essentially required for horizontal transmission and lifelong persistence, and has a poorly understood role in the development of viral malignancies (7). In addition to epithelial cells in the oropharynx (8), resting peripheral blood and tonsil memory B cells are thought to serve as reservoirs for latent EBV (2, 9), while mature B cell trafficking through the GC and terminal differentiation into CD38+ plasma cells can trigger EBV reactivation (10). In vitro, cross-linking of surface immunoglobulins (Ig) on freshly isolated EBV-positive B cells (11) or latently infected BL cells (12) functionally mimics antigen interactions and stimulates virus reactivation. The complex signaling events initiated through cross-linking of the B cell receptor (BCR) activate EBV immediate early (IE) genes BZLF1 and BRLF1, encoding viral transactivators Zta and Rta respectively, and induce the lytic cascade (7).

Exact molecular mechanisms, including underlying post-transcriptional processes, controlling EBV latency and the switch to lytic replication remain to be fully elucidated (7, 13). Master transcriptional regulators of plasma cell differentiation, such as Blimp-1/PRDM1, can activate the EBV Zta (Zp) and Rta (Rp) promoters (13, 14). Transcription factors such as ATF, Sp1/3, MEF2D, XBPs, CREB family members, AP1 heterodimers (i.e. c-Jun), and HIF1a interact with Zp in response to antigen stimulation or oxidative stress; Zp further contains cis-regulatory elements which confer auto-regulation (7, 13, 15–18). Repressors of Zp include the zinc finger E-box-binding proteins encoded by ZEB1 and ZEB2 and the polycomb protein Yin Yang 1 (YY1) (7, 13, 19–21). Notably, microRNAs (miRNAs) from the miR-200 family (miR-200b and miR-429 expressed in epithelial cells) post-transcriptionally silence ZEB1/2 expression, thereby modifying Zp activity (22, 23).

miRNAs are ~22 nucleotide (nt) non-coding RNAs that post-transcriptionally control gene expression and regulate multiple biological processes, including B cell development, GC reactions, and the progression of immune responses (24, 25). Deregulated miRNA activity is implicated in B cell lymphomagenesis, and normal B cell subtypes, as well as B cell cancers, can be distinguished by miRNA signatures (24, 26). EBV encodes >44 viral miRNAs, the majority of which are expressed from the BART locus, and exhibit expression kinetics similar to the BART transcripts (27). BART miRNAs are detectable throughout phases of EBV infection, including latency I (28), suggesting these molecules actively facilitate maintenance of the latent state (29, 30). However, viral miRNA expression is rapidly induced upon entry into the lytic cycle (27, 28), indicating a possible role in EBV reactivation. Recently, we identified several EBV miRNAs that attenuate BCR signal transduction and consequently, dampen BCR-induced EBV reactivation, demonstrating that a subset of the viral miRNAs actively suppress lytic replication (31).

The functions for most host miRNAs in EBV reactivation are not known. Subversion of apoptosis and evasion of anti-viral responses are key parts of viral reactivation, and several groups have previously demonstrated roles for miRNAs in these processes (30, 32). Additionally, studies in which Dicer was inhibited reported reduced levels of IE gene expression during EBV reactivation, providing evidence that components of the miRNA biogenesis machinery are necessary for aspects of lytic replication (33, 34). We previously reported that disruption of cellular miR-17 in EBV-positive BL cells augments IE gene expression (31), and other groups have shown that miR-200 family members expressed in epithelial cells have essential roles in the EBV latent to lytic switch (22, 23). In this study, we investigated how EBV exploits BCR-responsive cellular miRNAs to navigate entry into the lytic phase. Specifically, we aimed (i) to define the miRNAs that are altered by BCR-mediated lytic reactivation, (ii) to elucidate targets of those miRNAs, and (iii) to determine whether BCR-responsive miRNAs and/or their targets play a role lytic reactivation.

## Results

### BCR-mediated EBV reactivation induces changes in cellular miRNA levels

To investigate the role of host miRNAs in EBV reactivation, we treated EBV-positive MutuI cells with antibodies to surface Ig (aIgM) for 22 hrs and profiled the small RNAs by deep sequencing (<200 nt). Over one million sequences were obtained per sample, and following alignment to the human and EBV genomes, 1,015 distinct miRNAs (read count >1 in at least one library) were identified. Of these, 392 mature miRNAs, were considered expressed (read counts >=10) and used to determine differential expression (DE) in mock versus anti-IgM treated cells (Fig. 1A). Consistent with prior reports (27, 28), nearly all EBV miRNAs were upregulated following anti-IgM treatment (Fig. 1A). EBV miRNAs not shown (i.e. miR-BHRF1-2-5p) were still detected, but at levels below our cut-offs. In addition to viral miRNAs, we observed significant changes in host miRNAs. miR-141-3p, miR-146a-5p, miR-342-3p, miR-3609, miR-21-3p, and miR-21-5p increased upon BCR stimulation while 11 miRNAs, including miR-148a-5p, miR-27b-5p and miR-139-3p, significantly decreased.

**Figure 1.**
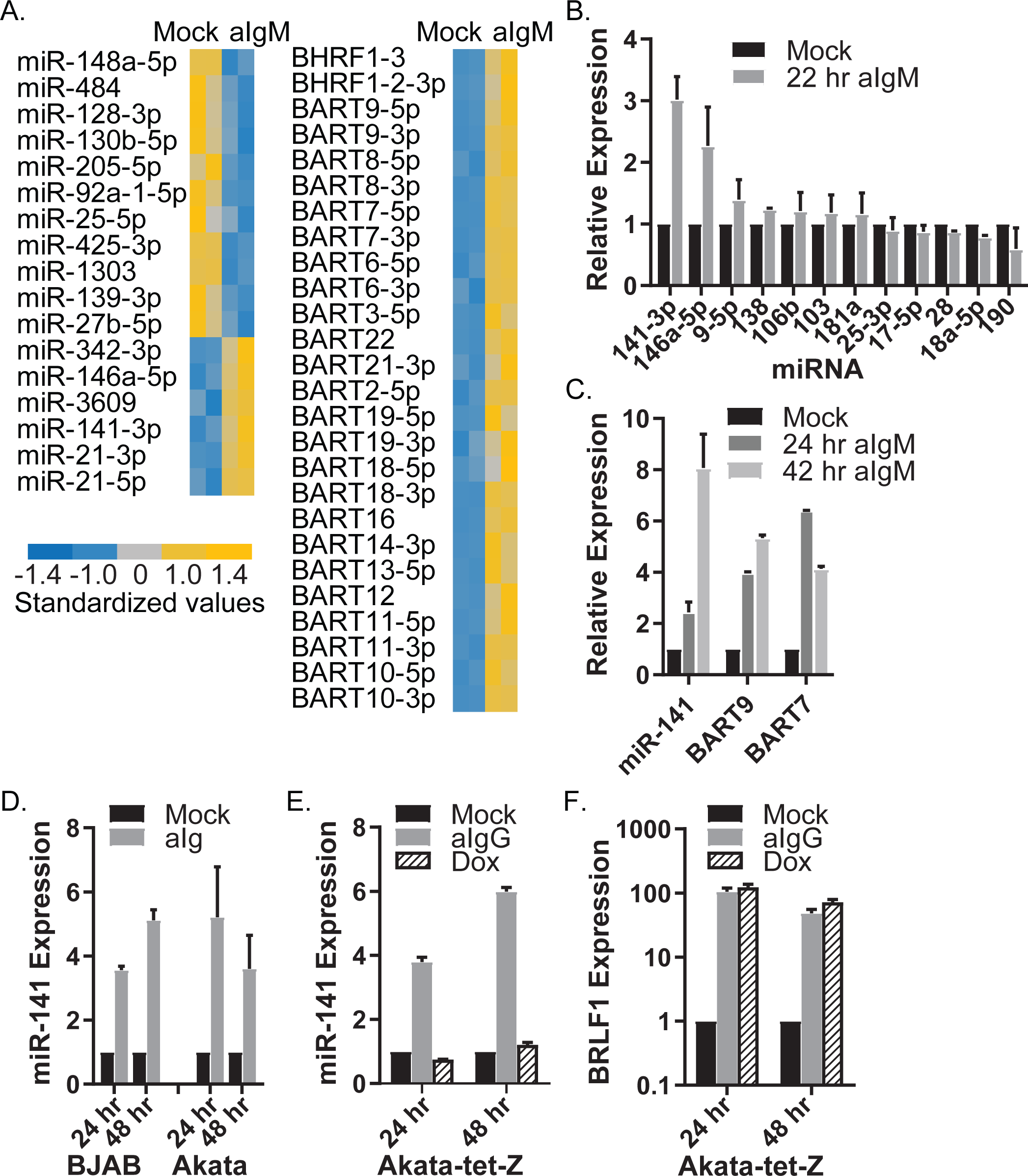
BCR cross-linking induces changes in the MutuI miRNAome. A. Heat-map of significantly D.E. EBV and cellular miRNAs in anti-IgM treated MutuI cells compared to mock (p<0.05, FDR<0.05, RC>20). Values for each miRNA are standardized across samples to a mean = 0 and standard deviation +/− 1. B. Taqman qRT-PCR for cellular miRNA expression. Total RNA was harvested from Mutu I cells treated with 2.5 ug/mL anti-IgM for 22 hrs. miRNAs were determined by qRT-PCR. Unless otherwise stated, in Figure 1, miRNA expression values are normalized to cellular miR-16 and reported relative to levels in mock cells. C. Taqman qRT-PCR analysis of miR-141, ebv-miR-BART9 and ebv-miR-BART7 expression in anti-IgM treated MutuI cells. D. and E. qRT-PCR analysis of miR-141 in anti-IgM treated EBV negative BJAB cells, anti-IgG treated EBV positive Akata cells, or anti-IgG or Dox treated Akata-tet-Z cells. F. Expression of EBV immediate-early (IE) gene BRLF1 in Akata-tet-Z cells following anti-IgG or Dox treatment. BRLF1 expression level was assayed by qRT-PCR. Values are normalized to GAPDH and shown relative to mock cells. Averages and standard deviations (S.D.) in Figure 1 are from at least three independent experiments.

BCR-mediated induction of miR-141-3p and miR-146a-5p were independently verified by qRT-PCR (Fig. 1B). We measured levels of other miRNAs which have been linked to BCR stimulation and/or EBV reactivation (i.e. miR-17/92, miR-181, miR-190) (37, 38), but did not detect robust changes in their expression (Fig. 1B). Enhanced miR-146a levels have been linked to LMP1 expression (39) while miR-21 is upregulated following EBNA2 expression (40) and de novo EBV infection (41). miR-342-3p is BCR-responsive in murine WEHI-231 cells (42). We therefore selected miR-141-3p for further analysis. While miR-141 has been studied extensively in epithelial cells, less is known about this miRNA in B cells. Similar to viral BART miRNAs, miR-141-3p accumulated during EBV reactivation (Fig. 1C).

### miR-141/200c are responsive to BCR cross-linking irrespective of EBV infection status

To investigate whether miR-141 induction is linked to any EBV factors, we tested additional BL cells. Surface Ig cross-linking increased miR-141-3p in both EBV-negative and EBV-positive cells (Fig. 1D), suggesting that miR-141 induction is not dependent upon EBV, but instead a cellular response to BCR signals. To definitively rule out EBV as a contributor, we tested miR-141-3p levels in EBV+ BL cells expressing a doxycycline inducible Zta (Akata-tet-Z) (43). In these cells, EBV reactivation can be initiated through the BCR or alternatively, through direct chemical induction of Zta. Treatment of Akata-tet-Z cells with anti-IgG led to increased miR-141-3p similar to what was observed for other BL cells; however, when EBV lytic replication was activated independent of the BCR, miR-141-3p levels were unaffected (Fig. 1E and F). We therefore conclude that miR-141-3p induction is linked to the normal cell signaling response initiated through BCR engagement and not mediated through any viral gene products.

miR-141 is part of the miR-200 family, consisting of five members that are processed from polycistronic primary miRNA transcripts (Fig. 2A) (44). While levels of other miR-200 family members were below our stringent cut-offs for miRNA profiling, we observed modest increases in miR-200c in response to anti-IgM (not shown), suggesting that the miR-141/200c pri-miRNA is transcribed. To explore this further, miR-200c, miR-429, and miR-200b levels were monitored in BL cells. Both miR-141-3p and miR-200c temporally accumulated with similar kinetics (Fig. 2B and C), consistent with activation of the pri-miRNA. In contrast, miR-429 and miR-200b were induced only in MutuI cells, indicating alternate modes of regulation (Fig. 2D and E).

**Figure 2.**
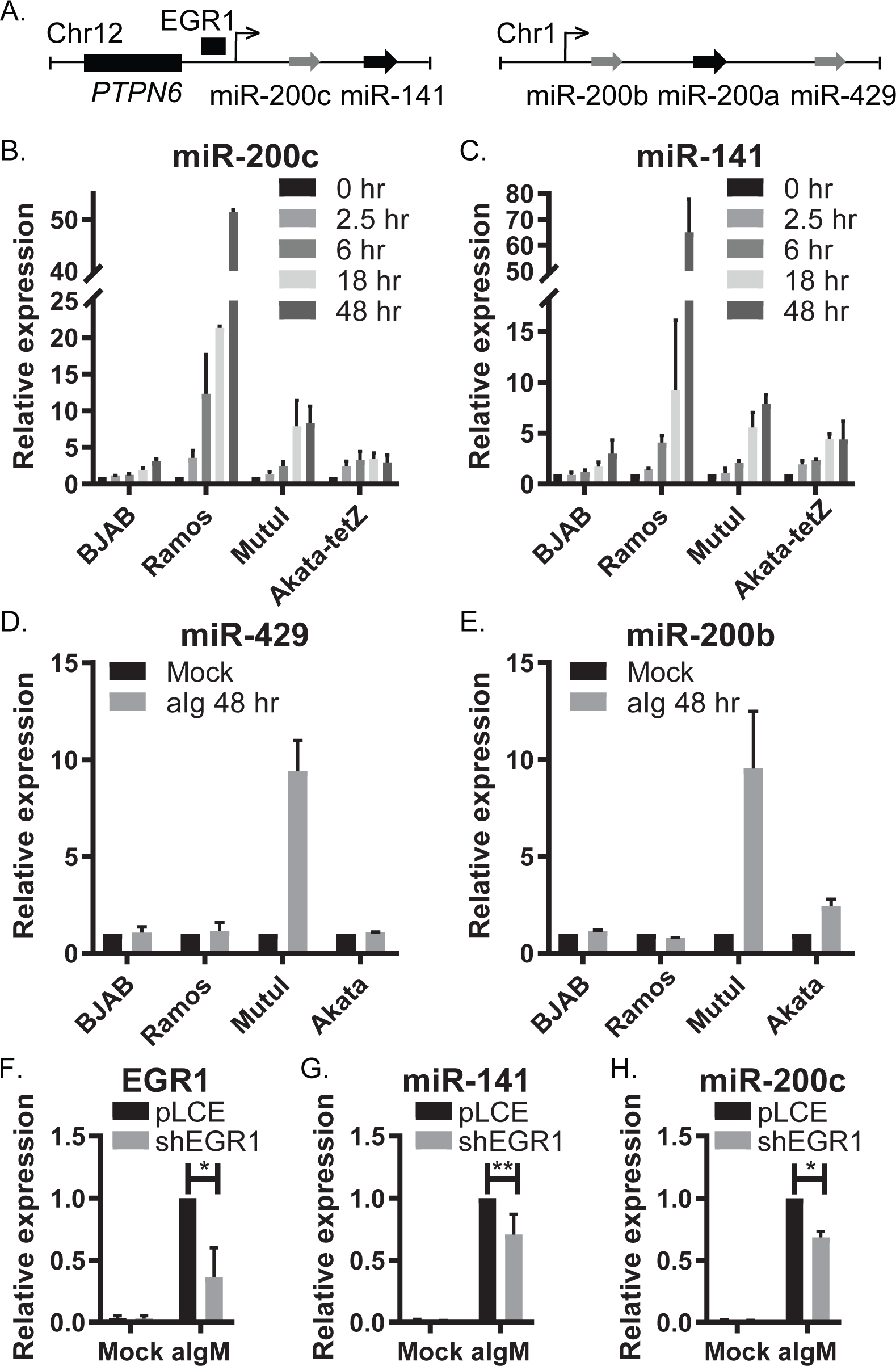
miR-141 and miR-200c, but not other miR-200 family members, are induced by BCR cross-linking. A. Schematic illustration of human chromosomes 12 and 1 harboring the gene loci of miR-200 family members, and their upstream regulatory modules. B. and C. qRT-PCR analysis of expression of miR-200c, miR-141 at various time points post-anti-Ig treatment of EBV-BJAB and Ramos cells and EBV+ MutuI and Akata-tet-Z cells. Values are normalized to miR-16 and reported relative to levels at 0 hr in each respective cell line. D. and E. qRT-PCR analysis of expression of miR-429 and miR-200b at 48 hrs post-anti-Ig treatment of the same panel of B cells. Values are normalized to miR-16 and reported relative to levels in each respective mock cell line. F. Knockdown of EGR1 in Ramos cells was assayed by qRT-PCR analysis. Expression levels are normalized to GAPDH and reported relative to anti-IgM treated control (pLCE) cells. G. and H. Induction of miR-141 and miR-200c by BCR cross-linking is impaired by EGR1 knockdown. miRNA expressions were analyzed by qRT-PCR. Values are normalized to miR-16 and reported relative to levels in anti-IgM treated control (pLCE) cells. Averages and S.D.s are from at least three independent experiments. By Student’s t-test, *p<0.05. **p<0.01.

The miR-141/200c promoter contains response elements for EGR1 (45), a rapid response nuclear zinc finger transcription factor that is upregulated upon BCR engagement (46). To explore EGR1 as a possible mechanism for how miR-141 and miR-200c are induced, we inhibited EGR1 with short hairpin RNAs (shRNAs) (Fig. 2F). Knockdown of EGR1 attenuated but did not fully abrogate miR-141 and miR-200c induction upon BCR cross-linking (Fig. 2G and 2H), pointing to a partial role for EGR1 in miR-141/200c transcription.

### miR-141 contributes to EBV reactivation

Prior studies in epithelial cells demonstrated positive correlations between EBV lytic gene expression and miR-200 family members (22, 23). To determine whether miR-141 plays a role in EBV reactivation in B cells, we utilized CRISPR-Cas9 to genetically inactivate miR-141 in MutuI cells expressing a tet-inducible Cas9 (iCas9). Following doxycycline treatment to activate Cas9, miRNA levels were measured by qRT-PCR, confirming specific disruption of miR-141 but not miR-200c (Fig. 3A and B). We subsequently examined EBV reactivation. Compared to control cells, disruption of miR-141 resulted in significantly reduced levels of the early EBV BMRF1 protein following surface Ig cross-linking (Fig. 3C and D). Consistent with attenuated lytic antigen expression, miR-141 inhibition also yielded decreases in viral loads and IE transcripts (Fig. 3E-G). Thus, miR-141 induction conferred through BCR signaling is necessary for efficient EBV reactivation.

**Figure 3.**
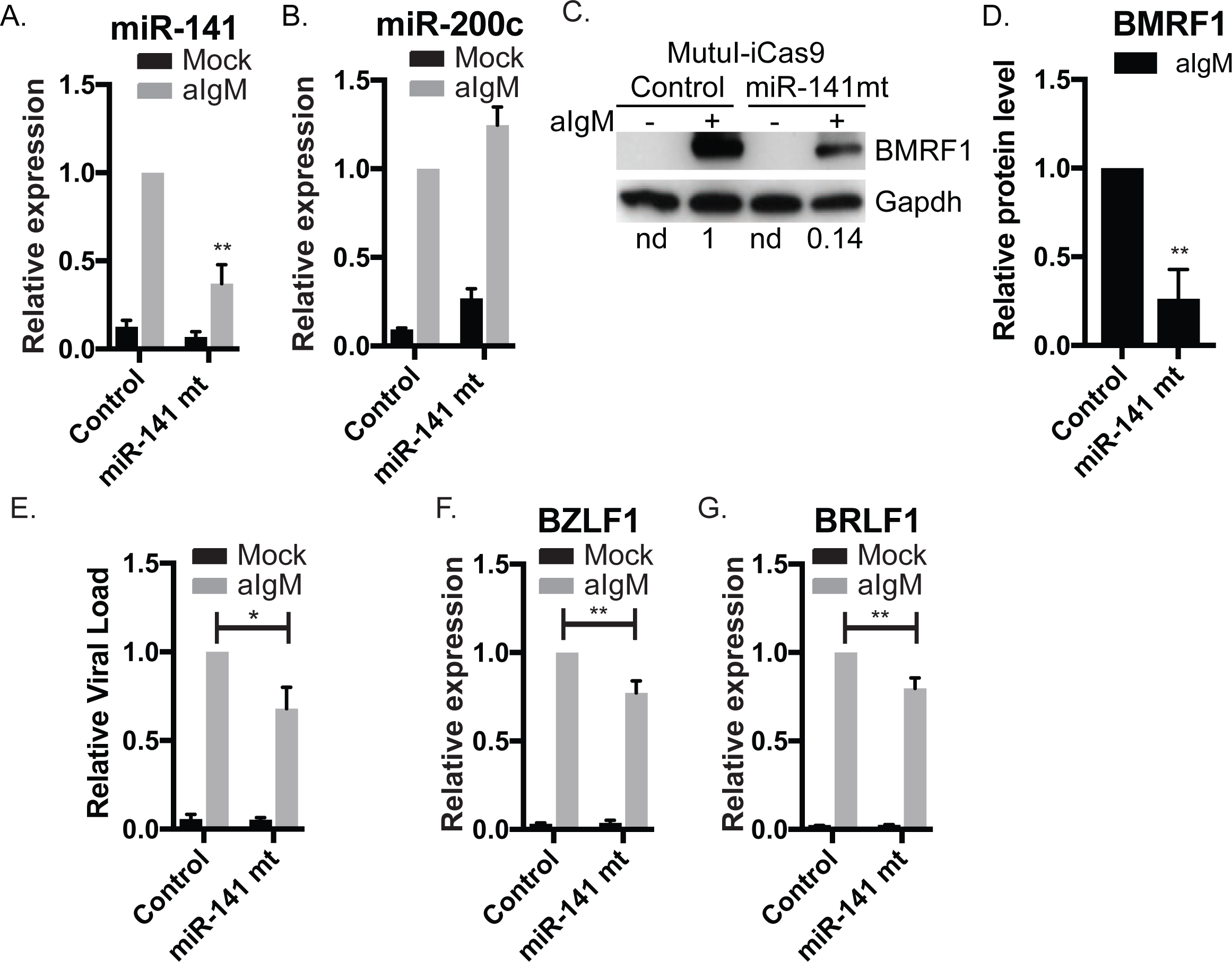
miR-141 contributes to efficient EBV lytic reactivation. A. and B. qRT-PCR analysis of miR-141 and miR-200c in MutuI-iCas9 confirms significant knockdown of miR-141 (A.) while miR-200c expression is not impaired (B.). Values are normalized to miR-16 and reported relative to levels of anti-IgM treated empty gRNA control cells. Student’s t-test, **p<0.01. C. Immunoblot of BMRF1 lytic gene product, Ea-D, in mock or anti-IgM treated MutuI-iCas9 cells stably transduced with either empty gRNA (Control) or gRNA against miR-141 (miR-141 mt). Gapdh levels are shown as loading controls. Band intensities were quantified using ImageJ, normalized to loading controls, and reported relative to anti-IgM treated empty gRNA control cells. Shown is the representative of four independent experiments. D. Quantification of immunoblot of MutuI-iCas9 for BMRF1 gene product. n=4. Student’s t-test, **p<0.01. E. Genomic DNA was isolated from mock or anti-IgM treated MutuI-iCas9 cells stably transduced with either empty gRNA or gRNA against miR-141. Viral loads were determined by qPCR assay using primers to the LMP1 region. Values are normalized to GAPDH and reported relative to viral levels in anti-IgM treated empty gRNA control cells. Shown is the average of four independent experiments. Student’s t-test, *p<0.05. F. and G. BZLF1 and BRLF1 expression levels were assayed by qRT-PCR. Values are normalized to GAPDH and shown relative to anti-IgM treated control cells. Shown are the averages of four independent experiments. Student’s t-test, **p<0.01.

### miR-141-3p and miR-BART9-3p function through common 5’ seed sequences

Intriguingly, EBV encodes a viral miRNA, miR-BART9-3p, which exhibits seed sequence homology to miR-141-3p (Fig. 4A), leading to the hypothesis that the viral and cellular miRNAs have common targets and common activities mediated through interactions with cognate seed match sites. To formally test this idea, miR-141 and miR-BART9 expression vectors were co-transfected with reporters harboring perfect binding sites in the luciferase 3’UTR for either miR-BART9-3p or miR-141-3p (Fig. 4B). As expected, both the viral and cellular miRNA potently downregulated their own reporters. Notably, we observed ~40% knockdown of luciferase activity when miR-141 was tested against the BART9 reporter and ~60% knockdown when miR-BART9 was tested against the miR-141 reporter (Fig. 4B). As the only stretch of sequence homology between these two miRNAs is the 5’ seed (nt 1-7), these results demonstrate that miR-141-3p and miR-BART9-3p are capable of functionally interacting with target RNAs through common 5’ sequences.

**Figure 4.**
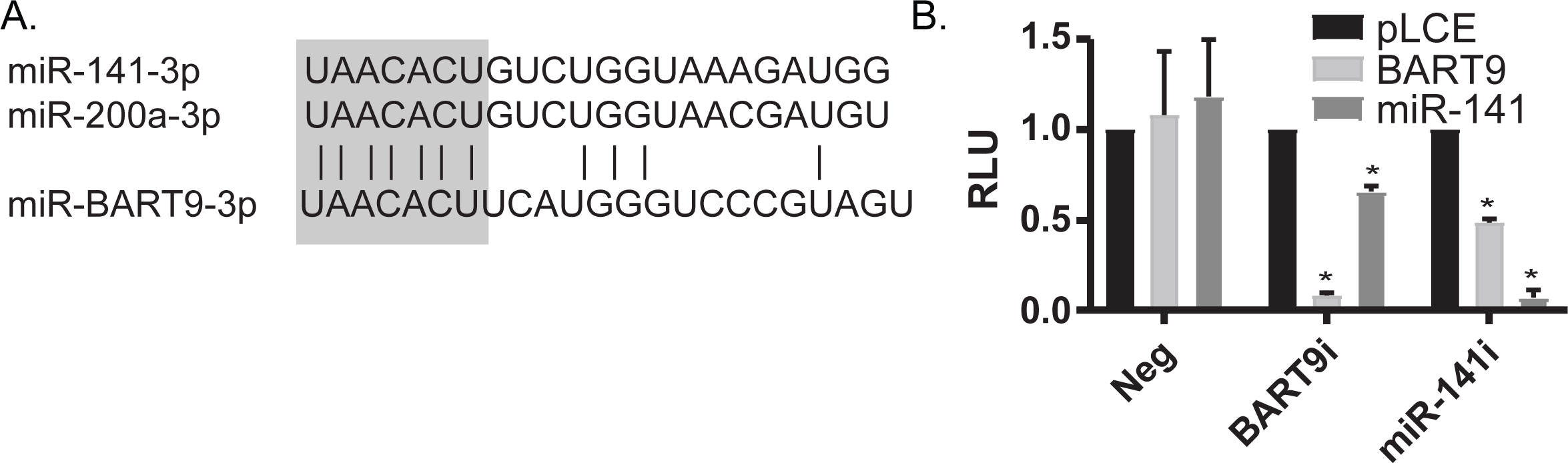
miR-141-3p and EBV miR-BART9-3p regulate 3’UTRs through common seed interactions. A. Alignment of miR-141-3p and miR-200a-3p with ebv-miR-BART9-3p. Sequence homology within the seed regions (nt 1-7) of these miRNAs is highlighted. B. miRNA expression vectors (pLCE or BART9 or miR-141) were co-transfected into HEK293T cells with reporters harboring perfect binding sites in the luciferase 3’UTR for either miR-BART9-3p (BART9i) or miR-141-3p (miR-141i). 48 hrs post-transfection, cells were lysed in 1X passive lysis buffer. Luciferase activity was measured using the dual luciferase reporter kit. Values are reported relative to empty reporter in control (pLCE) cells. Shown are the averages of three independent experiments performed in triplicate. Student’s t-test, *p<0.05. RLU=relative light units.

### Cellular targets are commonly regulated by miR-141 and miR-BART9

To investigate biological targets of miR-141-3p and miR-BART9-3p, we assembled a list of native 3’UTR interactions extracted from previously published studies (47–51). Target interactions were assigned based upon canonical miRNA seed pairing (i.e. nt 2-7). Additional candidates predicted by TargetScan (52) were included for a comprehensive list (Fig. 5A). We identified 1,880 unique 3’UTR targets harboring canonical 5’ seed matches to miR-141-3p, miR-200a, and/or miR-BART9-3p, of which 321 targets overlapped with one or more study. Comparison of the 187 3’UTRs captured in PAR-CLIP studies revealed that 41% of interaction sites could be assigned to both miR-BART9-3p and miR-141-3p based upon the presence of seed-match sites (inset, Fig. 5A).

**Figure 5.**
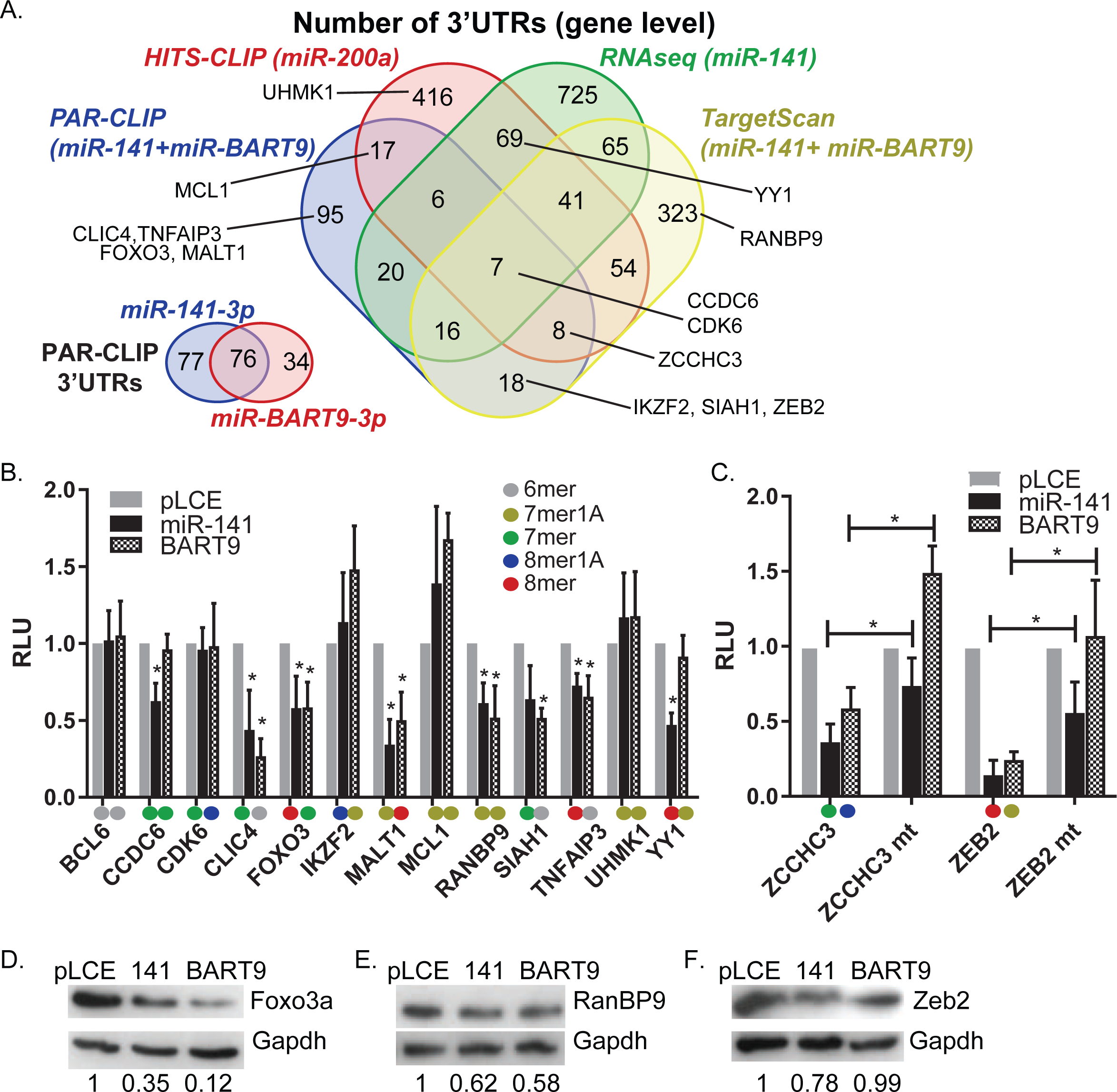
EBV miR-BART9-3p is a partial mimic of miR-141-3p. A. Overlap of miRNA interactions identified from published Ago-CLIP and RNAseq datasets and predicted by TargetScan. Reported are 3’UTR interactions with >7mer seed match to either miR-BART9-3p or miR-141-3p. B. Luciferase reporter assays confirm common 3’UTRs targeted by miR-141 and miR-BART9. HEK293T cells were co-transfected with psiCheck2 vector harboring selected 3’UTR luciferase reporters and EBV miRNA expression vectors (pLCE-based). 48-72 hrs post-transfection, cells were lysed and assayed for dual luciferase activity. Shown are the averages of at least three experiments performed in triplicate. Student’s t-test, *p<0.05. RLU=relative light units. The type of seed match between each individual miRNA and 3’UTR are labeled in different colors. C. miRNA binding sites in ZCCHC3 and ZEB2 3’UTRs were mutated by site-directed mutagenesis. Luciferase assays were performed as described in B. Shown are the averages of at least three experiments performed in triplicate. D-F. Immunoblots of Foxo3a, RanBP9, and Zeb2 in HEK cells transfected with miRNA expression vectors. Gapdh is shown as loading control.

miRNA interaction sites in CLIP and RNAseq studies are assigned computationally; thus, to experimentally confirm targets, luciferase reporters were constructed for 14 3’UTRs. We specifically selected 3’UTRs of interest to either B cell or EBV biology. These included (i) FOXO3, encoding Forkhead box O3a, a member of the conserved forkhead box transcription factors (53), (ii) RANBP9, encoding an adaptor protein that interacts with Zta to augment reactivation (54), (iii) YY1, necessary for GC reactions (55) and binds multiple sites within Zp to repress Zta expression (21), (iv) ZCCHC3, encoding a co-sensor for cGAS (56), and (v) ZEB2, a transcriptional repressor of Zp (13, 22). Binding sites in these 3’UTRs represented a range of 6mer (nt 2-7) to 8mer (nt 2-9) seed matches for miR-141-3p and/or miR-BART9-3p (Fig. 5B and C).

Reporters were tested in HEK293T cells co-transfected with miRNA expression vectors. For seven 3’UTRs, we observed significant inhibition of luciferase activity in the presence of either miRNA, supporting the idea that miR-BART9-3p functional mimics miR-141-3p (Fig. 5B and C). Mutagenesis of seed match sites in the ZCCHC3 and ZEB2 3’UTRs was sufficient to restore luciferase activity in the presence of EBV miR-BART9-3p and partially restored activity with miR-141 (Fig. 5C). Interestingly, luciferase assays further revealed distinct targeting of CCDC6 and YY1 3’UTRs by miR-141 but not miR-BART9 (Fig. 5B). Presumably, these targeting differences are due to 3’ compensatory binding.

We assessed the impact of miR-141 and miR-BART9 on endogenous Foxo3a, RanBP9, and Zeb2 protein levels in HEK293T cells. Foxo3a and RanBP9 were significantly reduced in response to both miRNAs (Fig. 5D and E). Surprisingly, despite the ZEB2 3’UTR responding to both miRNAs (Fig. 4D), we detected decreases in Zeb2 only in the presence of miR-141 (Fig. 5F), indicating that ZEB2 is not targeted by miR-BART9. Taken together, these experiments formally demonstrate that miR-BART9-3p acts as a partial mimic of miR-141-3p.

### miR-141 regulates Foxo3a levels in BL cells

Having demonstrated that Foxo3a levels are responsive to ectopic miR-141, we sought to determine whether FOXO3 was directly targeted by endogenous miR-141. EBV-negative Ramos-iCas9 and EBV-positive Akata-iCas9 cells were transduced with gRNAs to disrupt miR-141 (Fig 6B and E). Cells were treated with anti-Ig for 48 hrs and protein subsequently analyzed by immunoblotting. Compared to control cells, basal levels of Foxo3a increased by two to three fold in miR-141 mutant cells (Fig. 6A and D). miR-141 mutant cells were still responsive to BCR engagement; however, Foxo3a levels were consistently higher in these cells following anti-Ig treatment. We further tested viral copies in Akata-iCas9 cells by qPCR. Similar to observations in Fig. 3E, miR-141 disruption negatively impacted viral loads (Fig. 6G). Thus, perturbation of miR-141 in EBV-infected cells yields higher Foxo3a expression and is detrimental to lytic reactivation.

**Figure 6.**
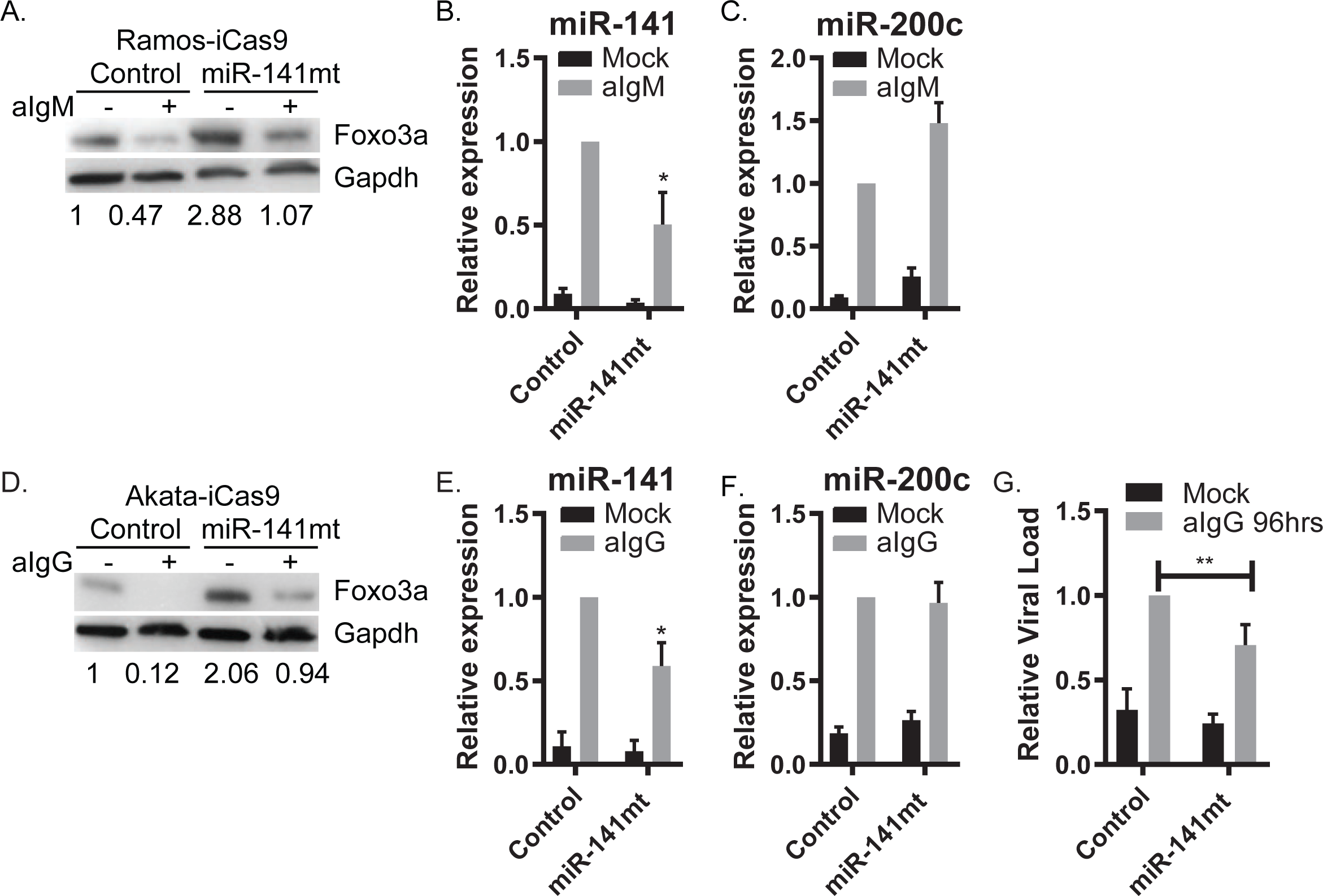
miR-141 regulates Foxo3a levels in BL cells. A. Immunoblot for Foxo3a in Ramos-iCas9 cells stably transduced with either empty gRNA (Control) or gRNA against miR-141 (miR-141 mt), indicating an enhanced level of Foxo3a when miR-141 knockdown is present. Gapdh levels are shown as loading controls. Band intensities were quantified using ImageJ, normalized to loading controls, and reported relative to mock treated empty gRNA control cells. Shown is the representative of two independent experiments. B. and C. qRT-PCR analysis of miR-141 and miR-200c in Ramos-iCas9 confirms knockdown of miR-141 (B.) while miR-200c expression is not impaired (C.). Values are normalized to miR-16 and reported relative to levels of anti-IgM treated empty gRNA control cells. Student’s t-test, *p<0.05. D. Immunoblot of Akata-iCas9 cells for Foxo3a. Shown is the representative of six independent experiments. Gapdh levels are shown as loading controls. Band intensities were quantified using ImageJ, normalized to loading controls, and reported relative to mock treated empty gRNA control cells. E. and F. qRT-PCR analysis of miR-141 and miR-200c in Akata-iCas9 confirms knockdown of miR-141 (E.) while miR-200c expression is not impacted (F.). Values are normalized to miR-16 and reported relative to levels of anti-IgG treated empty gRNA control cells. Student’s t-test, *p<0.05. G. Genomic DNA was isolated using DNAzol from mock or anti-IgG treated Akata-iCas9 cells stably transduced with either empty gRNA or gRNA against miR-141. Viral loads were determined by qPCR assay for LMP1 on genomic DNA. Values are normalized to GAPDH and reported relative to levels of anti-IgG treated empty gRNA control cells. Shown is the average of five independent experiments. Student’s t-test, **p<0.01.

### miR-BART9 enhances FOXO3 repression

To further understand the dynamics of Foxo3a repression during EBV reactivation, we monitored protein levels at multiple times post BCR stimulation (Fig. 7A and B). Foxo3a remained stable up until 24 hrs; by 48 hrs, however, total protein levels decreased three to five fold. Notably, this coincides with maximum miRNA induction (see Fig. 1). Similar to observations in Fig. 6, Foxo3a levels were lower in EBV-positive cells compared to EBV-negative cells, which is presumably due to miR-BART9 (Fig. 7C). Bypassing BCR signaling, Zta-mediated reactivation had little impact on Foxo3a, suggesting that viral gene products alone are not sufficient to measurably impact Foxo3a (Fig. 7C). Given that miR-141 is consistently induced in response to BCR engagement irrespective of infection status, we wondered whether miR-BART9 might exert an additive effect during infection. We therefore introduced miR-BART9 into EBV-negative cells (Fig. 7D). Ectopic miR-BART9 did not impact basal Foxo3a; however, upon BCR stimulation, miR-BART9 significantly enhanced Foxo3a reduction (Fig. 7E). These results indicate that the cellular environment conferred through BCR signaling, combined with the presence of miR-BART9, plays a key part in Foxo3a regulation during EBV reactivation.

**Figure 7.**
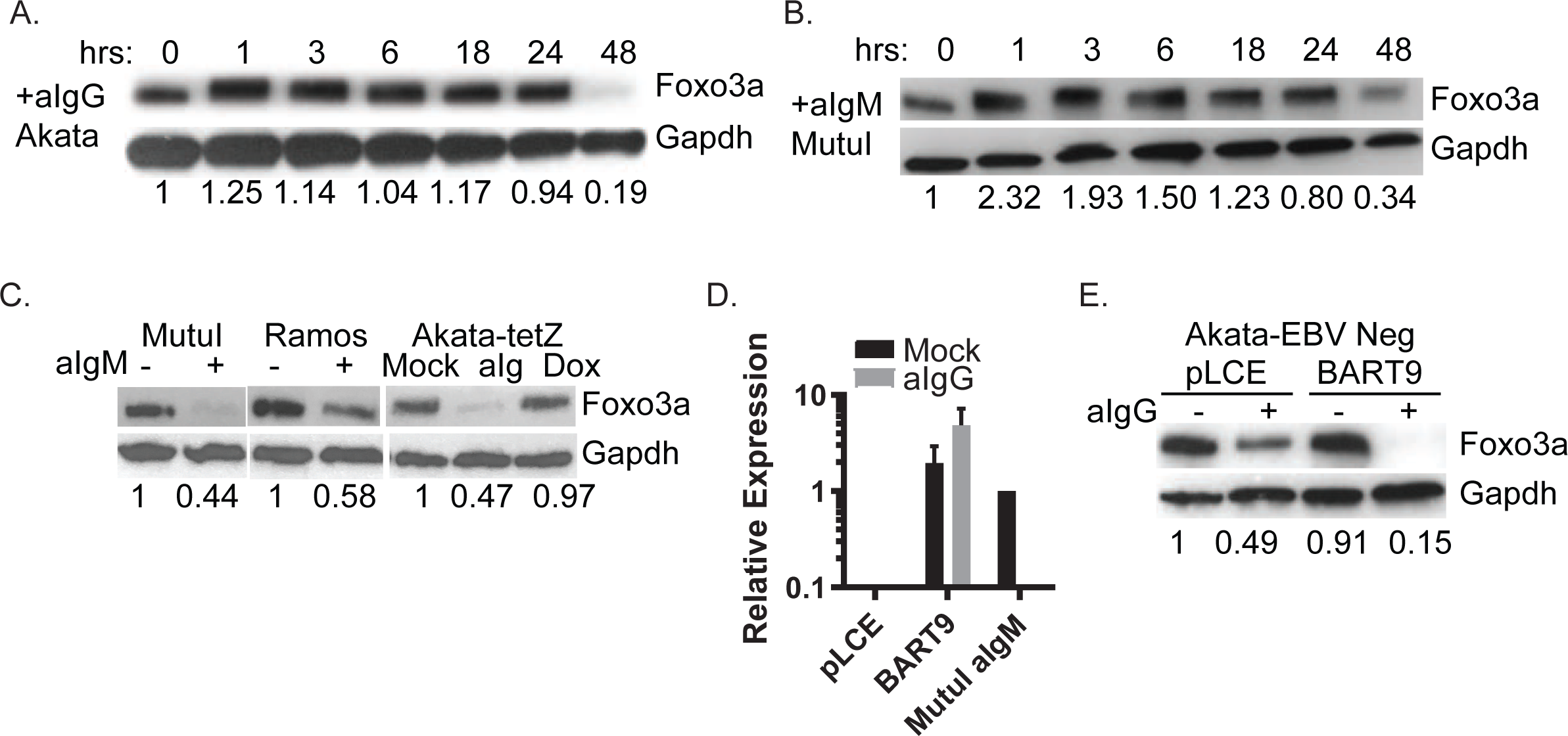
EBV miR-BART9 enhances Foxo3a inhibition. A. and B. Time course of Foxo3a protein levels in EBV-positive Akata cells treated with anti-IgG (A.) or MutuI treated with anti-IgM (B.). Significant reduction of total Foxo3a protein occurs between 24-48hrs. Shown is the representative of three independent experiments. C. BCR cross-linking but not Zta-mediated EBV reactivation reduces Foxo3a. Immunoblots were performed on lysates from selected B cells treated with anti-Ig to trigger BCR cross-linking or Akata-tetZ cells treated with Doxycycline to induce lytic reactivation. D. qRT-PCR analysis of miR-BART9 expression. EBV-negative Akata (Akata-EBV Neg) cells were transduced with pLCE-based miR-BART9 expression vectors. miR-BART9 expression values are normalized to cellular miR-16 and reported relative to levels in MutuI cells treated with anti-IgM for 48hrs. n=3. E. miR-BART9 enhances Foxo3a suppression following surface Ig cross-linking. Akata-EBV Neg cells transduced with pLCE-miR-BART9 were treated with anti-IgG five days post-transduction. After 48 hours of treatment, lysates were collected and subject to immunoblot as described above. Shown is the representative of three independent experiments.

### FOXO3 inhibition promotes EBV reactivation

Foxo3a levels inversely correlated with viral loads during reactivation (Fig. 6G), suggesting that Foxo3a might be restrictive for the lytic cycle. To test this hypothesis, we implemented shRNAs to post-transcriptionally block FOXO3 in EBV-positive cells (Fig. 8A and B). RNAi-mediated knockdown of FOXO3 sensitized cells to reactivation stimuli, and lead to significant increases in viral loads (Fig. 8A). We carried out similar experiments in EBV-positive Akata cells in which the lytic replication cycle can be activated directly through Zta. While miR-BART9 is expressed in these cells and accumulates in the presence of Zta, miR-141 is not induced in the absence of BCR stimulation, thereby allowing us to uncouple miR-141 and BCR signaling effects from EBV reactivation (Fig. 1 and 2). Compared to cells transduced with control vector, we observed significant increases in lytic antigen expression in shFOXO3 cells upon Zta expression (Fig. 8C and D). These findings demonstrate that the functional effects of BCR-mediated miR-141 induction can be phenocopied through direct inhibition of FOXO3. Moreover, via miR-BART9, EBV encodes counteractive measures to down-regulate expression of this cellular transcription factor which subsequently augments EBV reactivation.

**Figure 8.**
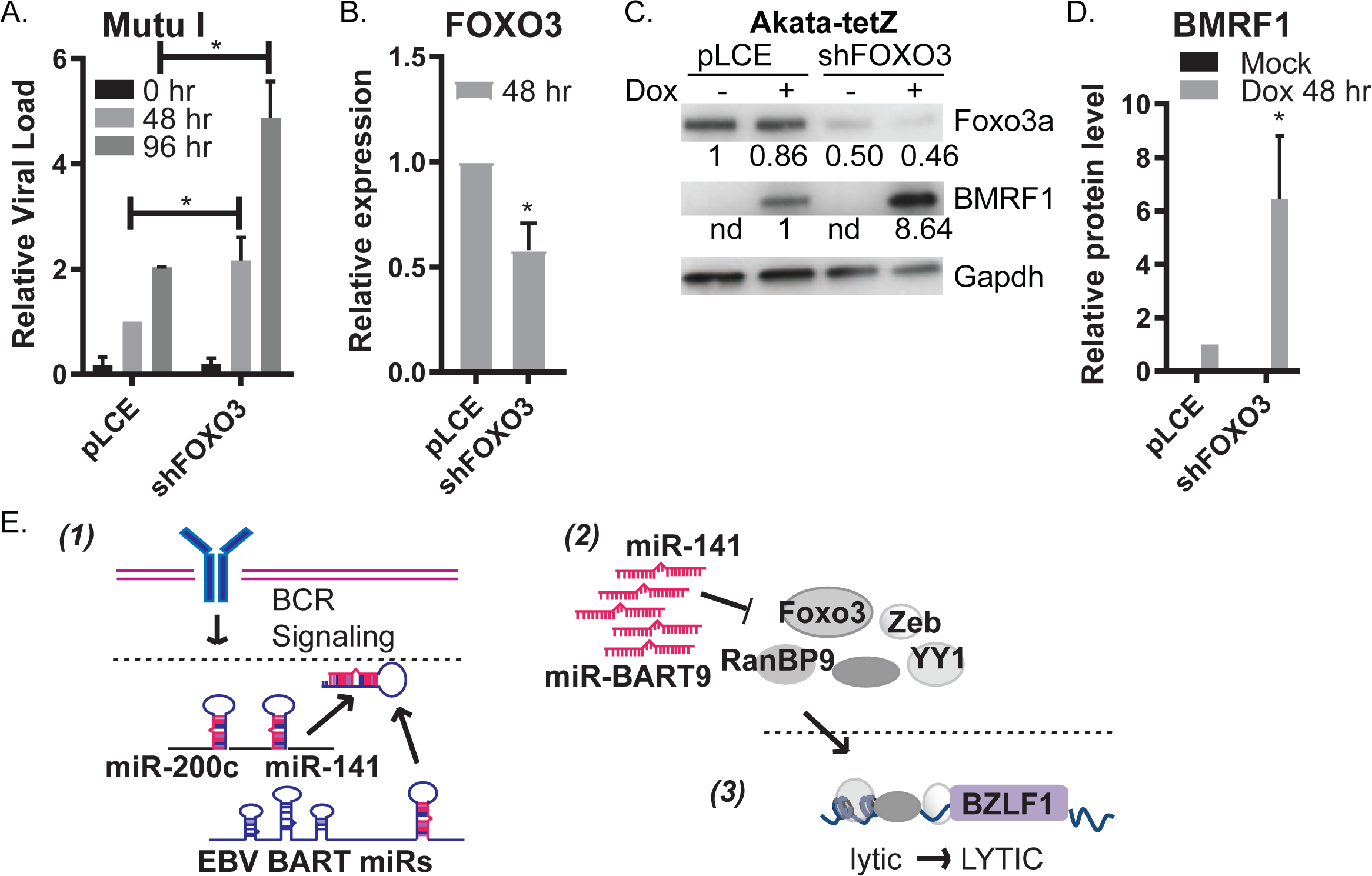
Inhibition of FOXO3 promotes EBV lytic reactivation. A. MutuI cells were stably transduced with empty vector control (pLCE) or shRNA against FOXO3 and treated with anti-IgM for 0, 48, or 96 hours. Genomic DNA was isolated using DNAzol and assayed by qPCR for LMP1. Values are normalized to GAPDH and reported relative to levels of 48hrs anti-IgM treated empty vector cells. Shown is the average of three independent experiments. Student’s t-test, *p<0.05. B. Confirmation of FOXO3 knockdown. Total RNA was collected from pLCE and shFOXO3 bearing MutuI cells treated with anti-IgM for 48hrs. FOXO3 expression was determined by qRT-PCR. Values are normalized to GAPDH and shown relative to anti-IgM treated control cells. Shown is the average of three independent experiments. Student’s t-test, *p<0.05. C and D. Akata-tetZ cells were stably transduced with empty vector control (pLCE) or shRNA against FOXO3 and treated with Doxycycline for 48hrs. Immunoblots were performed on lysates collected from mock or dox treated cells for Foxo3a and BMRF1 lytic gene product. Shown in C. is the representative of three independent experiments. Gapdh is shown as loading control. D. is the quantification of the immunoblots of BMRF1. Shown is the average of three independent experiments. Student’s t-test, *p<0.05. E. Proposed model of how BCR-responsive miR-141 and EBV miR-BART9 promote viral reactivation through suppression of common host targets that include FOXO3 and transcription factors known to associated with the Z promoter (YY1, ZEB1/2).

Taken together, our experiments support a model whereby BCR engagement triggers transcriptional activation of several miRNAs, including miR-141/200c (Fig. 8E). In EBV-infected cells, BCR engagement further activates lytic replication, leading to accumulation of viral miRNAs such as miR-BART9. Through seed-sequence mimicry, multiple targets of miR-141-3p are co-targeted by miR-BART9-3p, and while several identified targets (ZEBs, YY1, RANBP9) have previously described roles in repressing lytic reactivation, we demonstrate here that FOXO3 additionally restricts lytic reactivation. Thus, while miR-141 induction and FOXO3 suppression are physiological cellular responses to BCR triggers, EBV further harnesses and boosts this response through virally-encoded miR-BART9 to efficiently promote the lytic cycle.

## Discussion

In this study, we addressed the involvement of cellular miRNAs in EBV reactivation in B cells. Through in depth evaluation of miRNA expression, we found that miR-141, a member of the miR-200 family, was significantly induced in both EBV positive and negative BL cells in response to BCR stimulation. Interestingly, genetic disruption of miR-141 detrimentally impacted virus reactivation, suggesting that miR-141 and the direct targets of this miRNA are key factors in the EBV lytic replication cycle.

The functions of miR-200 family members in B cell processes, and specifically, EBV replication in B cells, are not fully known, which may be due in part to their complex expression patterns in B cell subsets. Initial miRNA sequencing studies detected miR-200 family members in naïve and memory B cells, but not in GC or plasma cells (58). miR-200 family members were found to be increased in tonsillar B cells compared to normal, resting CD19+ B cells, and B lymphoma cells (AIDS-DLBCL as well as non-AIDS DLBCL) (59). Moreover, analysis of biopsy samples from a cohort of 83 DLBCL patients suggested loss of miR-200c was linked to poor prognosis (59). Recent studies showed that miR-200a, miR-200b, miR-200c, miR-429, and miR-141 are upregulated in CD77-B cells (centroblasts, centrocytes, and plasmablast) compared to naïve and centroblasts (60). Thus, the presence of miR-200 family members is tightly linked to specific stages of B cell differentiation. Our experiments imply that surface Ig expression and triggering of BCR-response pathways could mechanistically explain the different levels of miR-141 and miR-200c detected in these prior studies (Fig. 2).

Accumulating evidence indicates that miR-141 and other miR-200 family members play crucial roles in the replication of both DNA and RNA viruses. Picornaviruses, such as enterovirus 71, induce miR-141 and miR-200c through EGR1; miR-141-3p subsequently targets eIF4E to aid in host shut-off, thereby positively impacting productive virus replication (45). Through the use of shRNAs, we show here that EGR1 is also partially responsible for miR-141 induction in BL cells (Fig. 2). In other virus systems, miR-141 may be detrimental to virus replication. For example, introduction of artificial miR-141 mimics into HepG2 cells interferes with hepatitis B virus replication (61). For herpesviruses, human cytomegalovirus (HCMV) harbors binding sites for miR-200b/c and miR-429 in the 3’UTR of the immediate early gene UL122 (encoding IE2) and mutational inactivation of the miRNA binding site leads to increased viral loads (62). Thus, miR-200 family members negatively impact HCMV replication, potentially during acute infection or reactivation from latency, by interfering with expression of the lytic cascade. In contrast, for EBV infected epithelial cells, miR-200b/c and miR-429 have been shown to suppress transcriptional repressors (ZEBs) of the EBV IE promoters, thereby indirectly enhancing lytic gene expression (22, 23). We found that miR-141 could target the ZEB2 3’UTR and reduce Zeb2 levels in HEK293T cells (Fig. 5). However, consistent with several studies examining Zeb expression in lymphocytes and other immune cell populations (63), we were unable to confirm Zeb2 expression in BL cells, thus prompting us to explore roles for other miR-141 targets in EBV reactivation.

Target identification for miR-141 in EBV-infected cells is complicated by the fact that EBV encodes a viral miRNA (miR-BART9-3p) with seed sequence homology to miR-141-3p (Fig. 4). While it has long been hypothesized that this viral miRNA might act as a functional mimic of miR-141 and/or other miR-200 family members, we formally demonstrate in this study that this is indeed the case. Using luciferase reporter assays, we show that multiple cellular 3’UTRs can be commonly targeted by both miR-141 and miR-BART9 (Fig. 5). Additionally, RanBP9 and Foxo3a protein levels were suppressed in the presence of either miRNA. Intriguingly, we identified two 3’UTRs (YY1 and CCDC6) that responded only to miR-141, despite strong seed matches to miR-BART9-3p. Furthermore, Zeb2 levels were inhibited by miR-141 but not miR-BART9, despite the 3’UTR reporter responding to both miRNAs (Fig. 5). While it remains to be specifically tested, these data strongly suggest that base-pairing outside of the seed region dictates the exact repertoire of targets for each miRNA and also contributes to the overall level of translational repression for each of their targets.

Given the potent inhibition of Foxo3a levels upon miR-141 or miR-BART9 expression, we selected this target as a potential candidate for involvement in the EBV latent-to-lytic switch. Foxo transcription factors have a wide range of functions and participate in a multitude of cellular processes including cell cycle, apoptosis, stress response, and cell differentiation. Notably, chemical inhibition of one member of the Foxo family, Foxo1, induces reactivation of the related human g-herpesvirus, KSHV, in adherent iSLK cells (64), suggesting an important role for Foxo proteins in modulating the herpesvirus latent-to-lytic switch. Studies in murine models have shown that PI3K/Akt/Foxo signaling contributes to multiple aspects of lymphocyte development including control of GC transcriptional programs and regulation of AID in activated B cells (65, 66). The majority of studies to date have focused on FOXO1, which is induced by EBF1, TCF3, and other B cell transcription factors, and plays an essential role in pro- and pre- B cell development (reviewed in (53)).

An understanding of the role of FOXO3 in B cell functions and EBV biology remains incomplete. FOXO3 transcripts are potently induced in human B cells committed to plasma cell differentiation (67). Post-translational modifications such as phosphorylation, acetylation, and glycosylation control subcellular localization and transcriptional activity of Foxo proteins (68). Cytoplasmic, phosphorylated Foxo3a is marked for degradation by the ubiquitylation-mediated proteasome degradation pathway while interactions between Foxo3a and 14-3-3 proteins result in nuclear shuttling (68). In the context of EBV infection, LMP1 can induce Akt activation and phosphorylation of Foxo3a, resulting in Foxo3a translocation out of the nucleus (69). Here, we demonstrate that the overall levels of Foxo3a are additionally controlled through both viral and host miRNA-mediated post-transcriptional regulation of FOXO3 transcripts. Through loss-of-function experiments, we further establish that FOXO3 inhibition positively impacts EBV reactivation (Fig. 8). Additional studies are needed to understand exact molecular functions of FOXO3 during this critical transition from latency to lytic replication.

Interestingly, in other virus infection systems, FOXO3 suppression is linked to anti-viral responses. FOXO3 blocks transcription of IRF7 in macrophages in response to poly-IC, and FOXO3 knockout mice exhibit increase lung injury during vesicular stomatitis virus (VSV) infection (70). Type I interferon can suppress FOXO3 levels (70) and more recent studies show this occurs partly through activation of miR-223 which targets the FOXO3 3’UTR (71). Of note, we identified another common target of miR-141 and miR-BART9, ZCCHC3 (Fig. 5), that functions as a co-sensor for cGAS, activates the IFNbeta promoter, and contributes to innate immune responses directed against DNA virus infection in the cytosol (56). Overexpression of ZCCHC3 enhances herpes simplex virus 1 (HSV1) mediated activation of interferon response genes (56). Thus, regulation of FOXO3 and ZCCHC3 levels by miR-141 and miR-BART9 may play an important role in modulating innate immune responses activated during EBV replication.

In summary, these data enhance our current understanding of how host and viral miRNAs contribute to EBV reactivation. To date, the majority of studies have pointed to roles for miRNAs in supporting latent infection for herpesviruses (72–74). Against convention, we demonstrate that specific miRNAs can also act to promote lytic reactivation, lending support to the idea that these molecules actively orchestrate aspects of latency and reactivation to cooperatively facilitate viral persistence within a host. Future work is needed to understand the specific relationships between miR-141 and miR-BART9 co-regulated factors which can reveal novel therapeutic targets for EBV-associated lymphomas.

## Methods

### Cell culture

BL cell lines were maintained at 37°C in a 5% CO2-humidified atmosphere in RPMI-1640 supplemented with 10% fetal bovine serum (FBS) and 1% penicillin, streptomycin, and L-glutamine (P/S/G). MutuI cells originated from the laboratory of Dr. Erik Flemington. EBV negative Akata cells were provided by Dr. Renfeng Li. Akata-tetZ cells were maintained in tetracycline-free media and provided by Dr. JJ Miranda with permission from Dr. Alison Sinclair. HEK293T cells were maintained in high glucose DMEM supplemented with 10% FBS and 1% P/S/G. For preparation of lentiviruses, HEK293T cells were plated in 15-cm plates in complete media and transfected using Polyethylenimine (PEI) with 15 ug lentivector, 9 ug pDeltaR8.75 and 6 ug pMD2G. Media was changed to complete RPMI-1640 between 8 hrs and 16 hrs post-transfection. Lentiviral particles were harvested by sterile filtration of the supernatant using a 0.45 micron filter at 48 and 96 hrs post-transfection and used to transduce ~1 to 5 × 10^6 cells. For BCR cross-linking, BL cells were spun down and plated at 0.5 × 10^6 cells in fresh media containing soluble anti-IgM or anti-IgG (Sigma) at concentrations and times indicated in figure legends (2.5-5 ug/mL for 22-48 hrs).

### Plasmids

pLCE-based miRNA expression vectors contain ~200 nt of the pre-miRNA as previously described (47). Functional miRNA expression was confirmed by indicator assays as previously described (47); the miR-141 indicator is from Addgene (#67632). To generate 3’UTR luciferase reporters for BCL6, CDK6, IKZF2, MCL1, ZCCHC3, and ZEB2, regions were PCR amplified from genomic DNA of EBV-infected B cells and cloned into the XhoI and NotI sites downstream of *Renilla* luciferase in the psiCheck2 dual luciferase reporter vector containing an expanded multiple cloning site (75). psiCheck2 3’UTR constructs for FOXO3, RANBP9, and YY1 were provided by Dr. Jay Nelson’s laboratory at VGTI. Additional 3’UTR reporters are cloned into pLSG as previously described (31, 47). Oligonucleotide sequences used for cloning are available upon request. Mutant 3’UTR reporters, containing nucleotide changes in miRNA seed match sites as identified by PAR-CLIP, were generated by Phusion Taq site-directed mutagenesis as previously described (76).

### CRISPR editing

Inducible Cas9 (iCas9) BL cells were established by transducing cells with pCW-Cas9-Blast-based lentiviruses (Addgene #83481) and selecting with Blasticidin. iCas9 cells were subsequently transduced with LentiGuide-puro lentiviral vector (Addgene #52963) bearing either empty guide RNA (gRNA) as control or gRNA against miR-141 and then selected with puromycin. Stable cell lines were treated with doxycycline for 7 days prior to analysis. Cells were treated with anti-IgM for 48 hours and lysates harvested for immunoblot analysis.

### Quantitative RT-PCR (reverse transcription-polymerase chain reaction) and PCR analysis

For gene expression analysis, total RNA was extracted using TRIzol (Thermofisher), DNAse-treated, and reversed transcribed using MultiScribe (Thermofisher) with random hexamers. Cellular and viral genes were detected using PowerUp SYBR Green qPCR (Thermofisher). Oligonucleotides sequences are available upon request. For viral loads, genomic DNA was isolated using DNAzol (Thermofisher). 100 ng of DNA were analyzed using primers to the LMP1 region and normalized to GAPDH levels as previously described (31). All PCR reactions were performed in technical replicates (duplicates or triplicates).

### 3’UTR reporter assays

HEK293T cells plated in 96-well black-well plates were co-transfected with 20 ng of 3’UTR reporter and 250 ng of control vector (pLCE) or miRNA expression vector using Lipofectamine2000 (Thermofisher). 48-72 hrs post-transfection, cells were harvested in 1X passive lysis buffer (Promega), and lysates were assayed for dual luciferase activity using the Dual Luciferase Reporter Assay System (Promega) and a luminometer. All values are reported as relative light units (RLU) relative to luciferase internal control and normalized to pLCE control vector.

### Western blotting

Cells were lysed in NP40-lysis buffer (50 mM HEPES pH 7.5, 150 mM KCl, 2 mM EDTA, 1 mM NaF, 0.5% (vol/vol) NP40, 0.5 mM dithiothreitol (DTT)). Protein concentrations were determined using the bicinchoninic acid (BCA) protein assay kit (Thermo Scientific), and 20 ug of total protein lysate were resolved on 10% Tris-glycine SDS-PAGE and transferred onto Immobilon PVDF membranes. Blots were probed with primary antibodies to Zeb2 (sc-271984, Santa Cruz), Foxo3a (2497S, Cell Signaling), RanBP9 (17755-1-AP, Proteintech), or Gapdh (sc-47724, Santa Cruz), followed by horse-radish peroxidase conjugated secondary antibodies (anti-rabbit IgG or anti-mouse IgG). Blots were developed with enhanced chemiluminescent substrate (Pierce). Band intensities were quantified using ImageJ, normalized to loading controls, and reported relative to control cells.

### miRNA deep sequencing and bioinformatics

miR-Seq libraries were generated from total RNA using the Illumina small RNA TruSeq kit as per manufacturer’s recommendations and sequenced multiplexed on the Illumina MiSeq at the ONPRC Molecular Biology Core. Prior to library preparation, purity of input RNA was assessed using the Nanodrop 2000 spectophotometer (ThermoScientific) and OD260/280 ratios of 1.8 to 2.1 were considered acceptable. Raw sequencing reads obtained in FASTQ format were preprocessed to remove linkers and aligned concurrently to the human genome (hg19) and Mutu I EBV genome (KC207814.1) using Bowtie (v1.0.1 http://bowtie-bio.sourceforge.net/index.shtml) (-v 2 –m 10) (77)). miRNAs were annotated and quantified by miRDeep (78). EdgeR (79) was used to define the significant, differentially expressed miRNAs in control versus anti-IgM treated cells (p<0.05, FDR<0.05, read counts >20). Heatmaps were generated in R.

Raw miR-Seq data files can be accessed through NCBI short read archive (SRA) (SUB6521585, submission in progress).

### Statistical analyses

Luciferase and PCR data are reported as mean of at least three independent experiments (unless otherwise stated) with standard deviations (S.D.). Statistical significance was determined by paired Student’s t test, performed using Microsoft Excel 2010, and values p < 0.05 were considered significant.

## Acknowledgements

This work was supported by a Pathway to Independence Award CA175181 from the National Cancer Institute, by R01 AI143620 from the National Institute of Allergy and Infectious Diseases, and by a pilot award from the Collins Medical Trust to RLS. The authors thank Isabella Brink, Robert Manni, and Camille Skinner, all previous undergraduate students in the Skalsky lab, for assistance with molecular cloning, and Yibing Jia at the ONPRC Molecular Biology Core (supported by P51 OD011092) for assistance with Illumina sequencing.

